## Supplementary figures S1-S3 for "A minimal model for microbial biodiversity can reproduce experimentally observed ecological patterns"

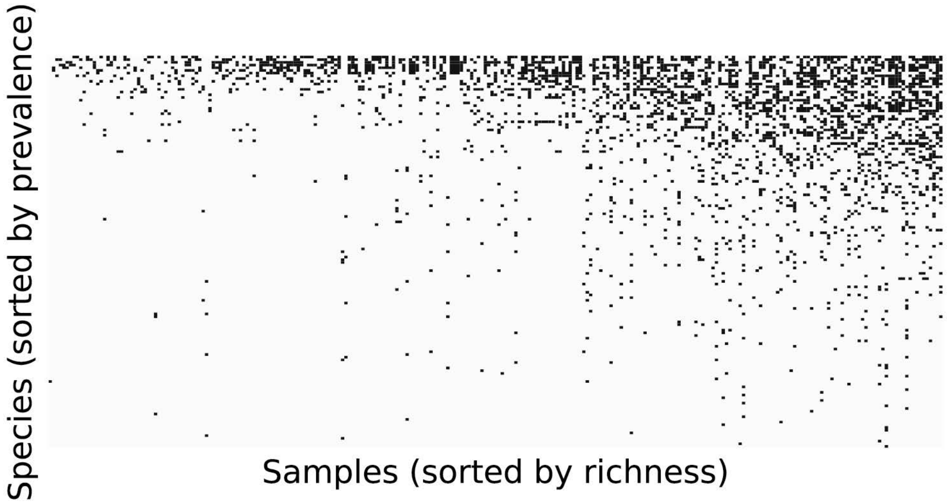


**Supplementary Figure S1.** Presence (black) and absence (white) of each species in simulations where both the environmental harshness m_env and the initial number of species varied from sample to sample (with sampling of both quantities following the same protocols as in Figure 2 of the main text).

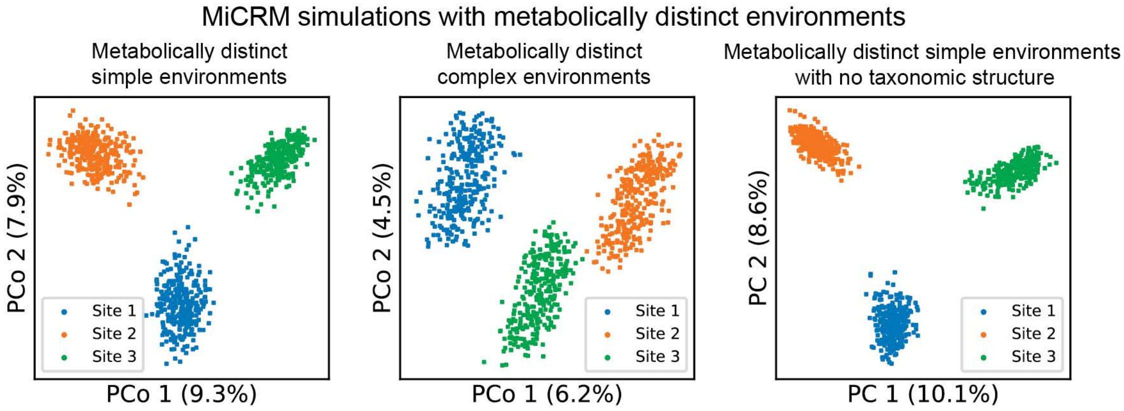


**Supplementary Figure S2**. PCoA of species-level compositions for the three simulated scenarios in which different body sites were supplied with resources from different classes. See Methods for details.

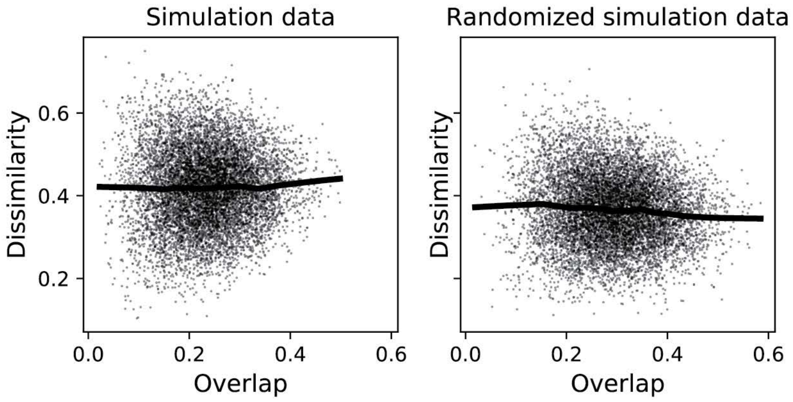


**Supplementary Figure S3.** Dissimilarity vs. overlap for pairs of samples taken from distinct body sites in the simulated HMP data.
